# Distinct conformational states of SARS-CoV-2 spike protein

**DOI:** 10.1101/2020.05.16.099317

**Authors:** Yongfei Cai, Jun Zhang, Tianshu Xiao, Hanqin Peng, Sarah M. Sterling, Richard M. Walsh, Shaun Rawson, Sophia Rits-Volloch, Bing Chen

## Abstract

The ongoing SARS-CoV-2 (severe acute respiratory syndrome coronavirus 2) pandemic has created urgent needs for intervention strategies to control the crisis. The spike (S) protein of the virus forms a trimer and catalyzes fusion between viral and target cell membranes - the first key step of viral infection. Here we report two cryo-EM structures, both derived from a single preparation of the full-length S protein, representing the prefusion (3.1Å resolution) and postfusion (3.3Å resolution) conformations, respectively. The spontaneous structural transition to the postfusion state under mild conditions is independent of target cells. The prefusion trimer forms a tightly packed structure with three receptor-binding domains clamped down by a segment adjacent to the fusion peptide, significantly different from recently published structures of a stabilized S ectodomain trimer. The postfusion conformation is a rigid tower-like trimer, but decorated by N-linked glycans along its long axis with almost even spacing, suggesting possible involvement in a mechanism protecting the virus from host immune responses and harsh external conditions. These findings advance our understanding of how SARS-CoV-2 enters a host cell and may guide development of vaccines and therapeutics.

## Introduction

The current coronavirus outbreak has become a pandemic reaching nearly every country on the planet, with a high case-fatality rate and devastating social and economic consequences. Coronaviruses (CoVs) are enveloped positive-stranded RNA viruses, including the two that caused previous outbreaks of severe acute respiratory syndrome (SARS) and Middle East respiratory syndrome (MERS), both with significant fatalities^1-3^, as well as several endemic common-cold viruses^4^. With a large number of similar viruses circulating in bats and camels^5-8^, experts have repeatedly warned that additional outbreaks are inevitable and pose major threats to global public health. The disease caused by a new virus SARS-CoV-2^9^, now named COVID19 (coronavirus disease 2019) by WHO, has created urgent needs for diagnostics, therapeutics and vaccines to contain the ongoing crisis, as well as potential future needs should it become seasonal. Meeting these needs requires a deep understanding of the structure-function relationships of viral proteins and relevant host factors.

For all enveloped viruses, membrane fusion is a key early step for entering host cells and establishing infection^10^. Although an energetically favorable process, membrane fusion has high kinetic barriers when two membranes approach each other, mainly due to repulsive hydration forces^11,12^. For viral membrane fusion, free energy to overcome these kinetic barriers comes from refolding of virus-encoded fusion proteins from a primed, metastable prefusion conformational state to a stable, postfusion state^13-15^. The fusion protein for CoV is its spike (S) protein that decorates the virion surface as an extensive crown (hence, “corona”). The protein also induces neutralizing antibody responses and is therefore an important target for vaccine development^16^. The S protein is a heavily glycosylated type I membrane protein anchored in the viral membrane. It is first produced as a precursor that trimerizes and possibly undergoes cleavage, potentially by a furin-like protease, into two fragments: the receptor-binding fragment S1 and the fusion fragment S2 (Fig. 1A; ref^17^). Binding through the receptor-binding domain (RBD) in S1 to a host cell receptor (e.g., angiotensin converting enzyme 2 (ACE2) for both SARS-CoV and SARS-CoV-2) and further proteolytic cleavage at a second site in S2 (S2’ site), by a serine protease TMPRSS2^18^ or endosomal cysteine proteases cathepsins B and L (CatB/L), are believed to trigger possible dissociation of S1 and irreversible refolding of S2 into a postfusion conformation – a trimeric hairpin structure formed by heptad repeat 1 (HR1) and heptad repeat 2 (HR2)^19,20^. These large structural rearrangements can bring together the viral and cellular membranes, ultimately leading to fusion of the two bilayers.

**Figure 1.**
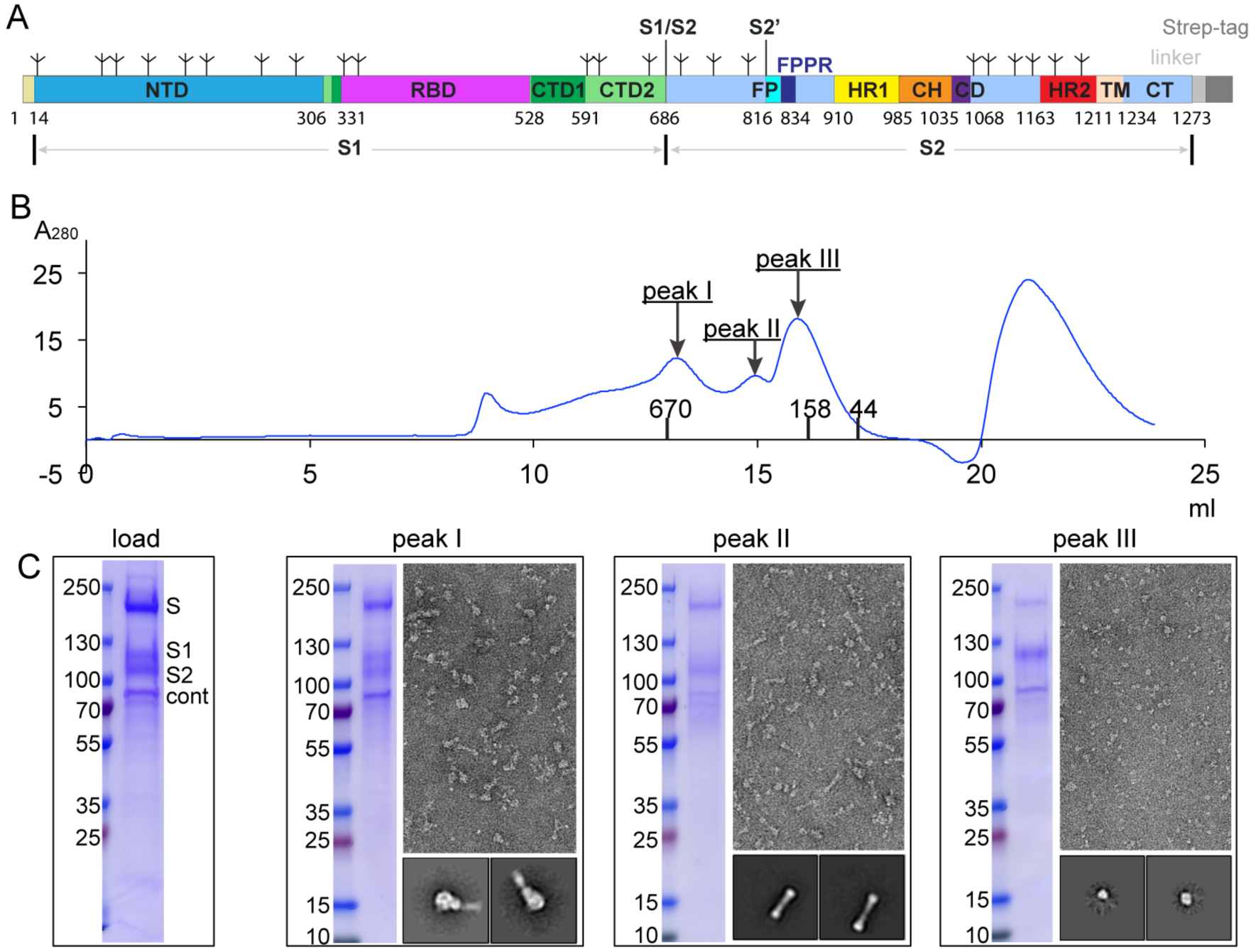
Preparation of a full-length SARS-CoV-2 spike protein. (A) Schematic representation of the expression construct of full-length SARS-CoV-2 spike (S) protein. Segments of S1 and S2 include: NTD, N-terminal domain; RBD, receptor-binding domain; CTD1, C-terminal domain 1; CTD2, C-terminal domain 2; S1/S2, S1/S2 cleavage site; S2’, S2’ cleavage site; FP, fusion peptide; FFPR, fusion peptide proximal region; HR1, heptad repeat 1; CH, central helix region; CD, connector domain; HR2, heptad repeat 2; TM, transmembrane anchor; CT, cytoplasmic tail; and tree-like symbols for glycans. A strep-tag was fused to the C-terminus of S protein by a flexible linker. (B) The purified S protein was resolved by gel-filtration chromatography on a Superose 6 column in the presence of detergent NP-40. The molecular weight standards include thyoglobulin (670 kDa), γ-globulin (158 kDa) and ovalbumin (44 kDa). Three major peaks (peak I-III) contain the S protein. (C) Load sample and peak fractions from (B) were analyzed by Coomassie stained SDS-PAGE. Labeled bands were confirmed by western blot (S, S1 and S2) or protein sequencing (S2 and Cont; S and S1 bands did not gave any meaningful results probably due to a blocked N-terminus). Cont, copurified contaminating protein, identified as endoplasmic reticulum chaperone BiP precursor by N-terminal sequencing. Representative images and 2D averages by negative stain EM of three peak fractions are also shown. The box size of 2D averages is ∼510Å.

Since the first genome sequence of SARS-CoV-2 was released^21^, several structures have been reported for S protein fragments, including its ectodomain stabilized in the prefusion conformation^22,23^ and RBD-ACE2 complexes^24-26^ (Fig. S1), largely building upon the previous success of the structural biology of S proteins from other CoVs^20^. The stabilized S ectodomain of the new virus adopts an architecture almost identical to that of the SARS-CoV ectodomain, with S1 folding into four domains - NTD (N-terminal domain), RBD, and two CTDs (C-terminal domains) and protecting the prefusion conformation of S2 with HR1 bending back towards the viral membrane (Fig. S1A and S1B). The RBD samples two distinct conformations – “up” representing a receptor-accessible state and “down” representing a receptor-inaccessible state. The structures of RBD-ACE2 complexes show that the RBD folds around a five-stranded β sheet and presents a gently concave surface, which cradles the N-terminal lobe of ACE2, with a large interface mediated primarily by hydrophilic interactions (Fig. S1C and S1D). Other related structures representing the postfusion state of S2, exemplified by one derived from mouse hepatitis virus (Fig. S1E) and a lower-resolution one from SARS-CoV (Fig. S1F), suggest how the proposed structural rearrangements of S2 may proceed to promote membrane fusion and viral entry^27,28^. Comparison of the pre- and post-fusion states reveals that HR1 undergoes a “jack-knife” transition that can insert the fusion peptide (FP) into the target cell membrane. Folding back of HR2 places the FP and transmembrane (TM) segments at the same end of the molecule; this proximity causes the membranes with which they interact to bend toward each other, effectively leading to membrane fusion. In the previous structures, the regions near the viral membrane are either not present or disordered, and yet they all appear to play critical structural and functional roles^29-33^.

In the work reported here, we have expressed, in HEK293 cells, and purified a full-length, fully wild-type form of the SARS-CoV-2 S protein and determined two cryo-EM structures representing its prefusion and postfusion states, both derived from a single preparation solubilized in detergent. The prefusion structure is in the all-“down” configuration, and local differences from the structure of the soluble, stabilized ectodomain suggest that the latter may have antigenic properties that differ from those of the virion-borne spike. Spontaneous rearrangement to the postfusion conformation, documented here in the absence of any receptor, may also occur on the virion surface. We speculate that its presence could stabilize the virion during host-to-host transmission and could also have consequences for the antigenicity and immunogenity of the virion.

## Results

### Purification of intact S protein

To produce a functional SARS-CoV-2 S protein, we transfected HEK293 cells with an expression construct of a full-length wildtype S sequence with a C-terminal strep-tag (Fig. 1A). These cells fused efficiently with cells transfected with an intact human ACE2 construct, even without addition of any extra proteases (Fig. S2), suggesting that the S protein expressed on the cell surfaces is fully functional for membrane fusion. The fusion efficiency was not affected by the C-terminal strep-tag. To purify the full-length S protein, we lysed the cells and solubilized all membrane-bound proteins in detergent NP-40. The strep-tagged S protein was then captured on strep-tactin resin. The purified S protein eluted from a size-exclusion column as three distinct peaks (Fig. 1B). When the peak fractions were analyzed by Coomassie-stained SDS-PAGE (Fig. 1C), peak 1 contained both the uncleaved S precursor and the cleaved S1/S2 complex; peak 2 had primarily the cleaved but dissociated S2 fragment; and peak 3 included mainly the dissociated S1 fragment, as judged by N-terminal sequencing and western blot. Analysis by negative stain EM confirmed that particles resembling the prefusion S trimer dominated the peak 1 sample; those similar to the postfusion S2 were the major component of the peak 2 sample; and those in peak 3 had the size expected for a monomeric S1 (Fig. 1C). Protein from all three peaks showed binding to soluble ACE2 (Fig. S3), as each species was not well separated from the others by gel filtration chromatography, but peak 1 showed the strongest binding, comparable to that for the purified soluble S ectodomain trimer, while peak 2 showed the weakest binding, since it contained mainly the S2 fragment. Binding by protein from peak 3 was weaker than that of protein from peak 1, suggesting that the ability of monomeric S1 to bind ACE2 is somewhat weakened as compared to the S trimer. While the cleavage at the S1/S2 (furin) site is clearly demonstrated by protein sequencing of the N-terminus of the S2 fragment in peak 2, cleavage at the S2’ site is not obvious. We observed in some preparations a band around 20 kDa, a size expected for the S1/S2-S2’ fragment (Fig. 1C). We obtained a similar gel filtration profile when another detergent (DDM) was used to solubilize the S protein, although the peaks were less well resolved than the NP-40 preparation (Fig. S4), suggesting that the S protein dissociation is not triggered by any specific detergent. Finally, we also identified a major contaminating protein in the preparation as endoplasmic reticulum chaperone BiP precursor^34^, which may have a role in facilitating S protein folding.

### Cryo-EM structure determination

Cryo-EM images were acquired with selected grids prepared from all three peaks, on a Titan Krios electron microscope operated at 300 keV and equipped with a BioQuantum energy filter and a Gatan K3 direct electron detector. We used RELION^35^ for particle picking, two-dimensional (2D) classification, three dimensional (3D) classification and refinement. Structure determination was performed by rounds of 3D classification, refinement and masked local refinement, as described in Methods and Supplemental Materials. The final resolution was 3.1Å for the prefusion S protein; 3.3Å for the S2 in the postfusion conformation (Fig. S5-S8).

### Structure of the prefusion S trimer

The overall architecture of the full-length S protein in the prefusion conformation is very similar to the published structures of a soluble S trimer stabilized by a C-terminal foldon trimerization tag and two proline substitutions at the boundary between HR1 and the central helix (CH) (Fig. S1; ref^22,23^). Our new structure appears to be more stable, however, because the N-terminus, several peripheral loops and glycans are ordered (Fig. 2A and 2B), although invisible in those soluble trimer structures. As described previously, the S1 fragment has four domains, including NTD, RBD, CTD1 and CTD2 -- which all wrap around the three-fold axis, covering the S2 fragment underneath. Since the furin cleavage site at the S1/S2 boundary is in a surface-exposed and disordered loop (Fig. 2B), we do not know whether this structure represents the uncleaved or cleaved trimer, although the sample clearly contains the both forms (Fig. 1C). Nevertheless, the location of the furin site suggests that the cleavage may have little impact on the rest of the trimer structure, other than ultimately allowing complete dissociation of S1 fragment, which may be a key event that enables S2 refolding and membrane fusion. Likewise, the S2 fragment has a conformation nearly identical to that in the previous trimer structures, with most of the polypeptide chain packed around a central three-stranded coiled coil formed by CH, including the connector domain (CD), which links CH and the C-terminal HR2 through an additional linker region. Major differences between our new structure and the published trimer structures are that a ∼25-residue segment in S2 immediately downstream of the fusion peptide is ordered and S density at the C-terminus extends further to Pro1162 before fading away. The remaining segments, including HR2, TM and CT, are still not visible, at least in the high resolution maps.

**Figure 2.**
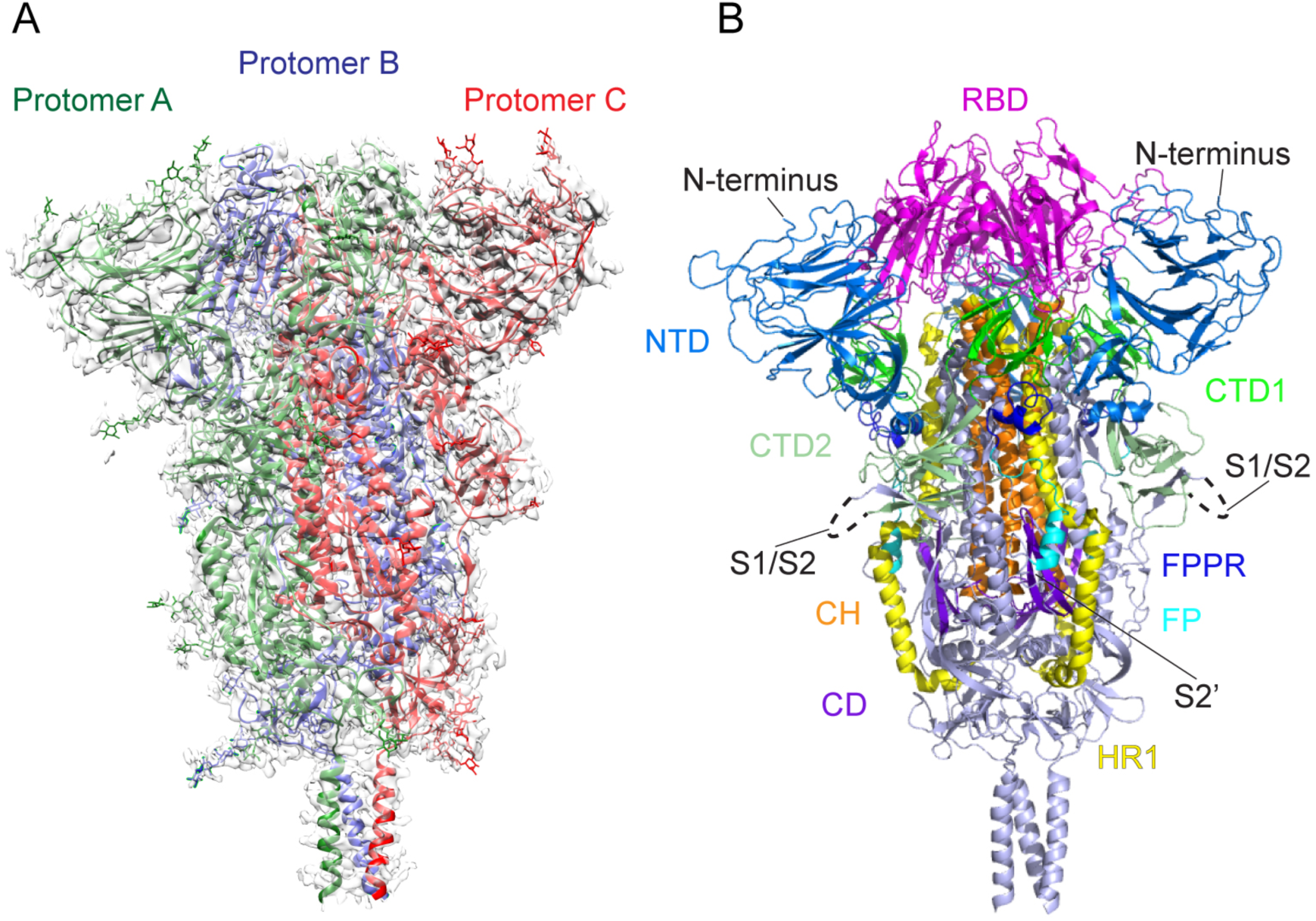
Cryo-EM structure of the SARS-CoV-2 S protein in the prefusion conformation. (A) The structure of the S trimer was modeled based on a 3.1Å density map. Three protomers (A, B, C) are colored in green, blue and red, respectively. (B) Overall structure of S protein in the prefusion conformation shown in ribbon representation. Various structural components in the color scheme shown in Fig. 1A include NTD, N-terminal domain; RBD, receptor-binding domain; CTD1, C-terminal domain 1; CTD2, C-terminal domain 2; FP, fusion peptide; FPPR, fusion peptide proximal region; HR1, heptad repeat 1; CH, central helix region; and CD, connector domain. N-terminus, S1/S2 cleavage site and S2’ cleavage site are indicated.

Several new features set our structure apart from the previously described prefusion conformations. First, the N-terminus in our new structure is ordered, including a disulfide bond (Cys15-Cys136) and a N-linked glycan at Asn17 (Fig. 3A), adopting a conformation similar to that of the N-terminus of the SARS-CoV S trimer^36^. It is unclear why this region is disordered in the published structures of the stabilized soluble S trimer from the new virus^22,23^, leaving an unpaired sulfhydryl group at Cys136. It would be important to confirm whether this region is folded in these constructs or simply poorly defined by density, despite a disulfide bond, particularly if they are widely used for vaccine studies.

**Figure 3.**
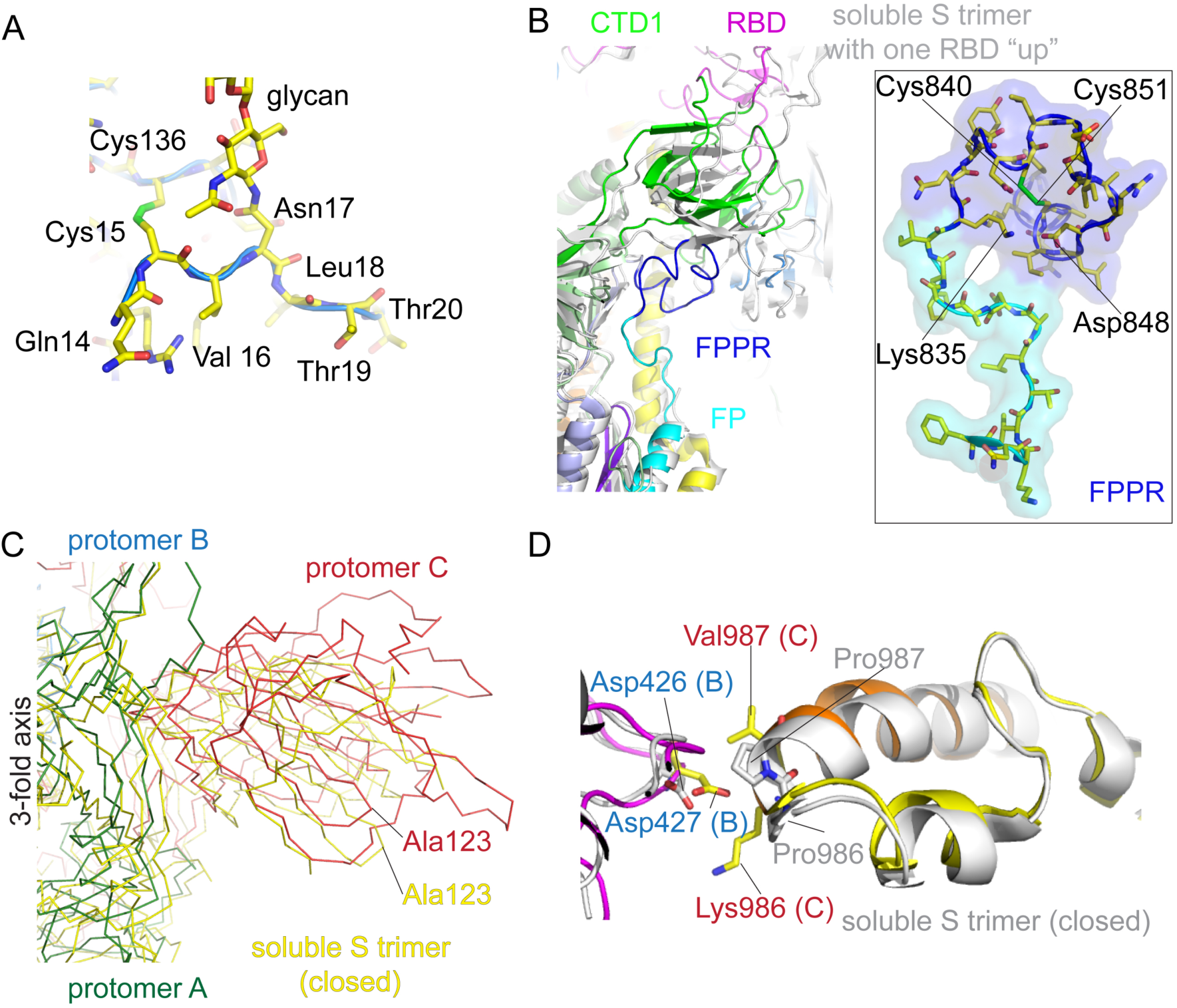
Selected new features of the SARS-CoV-2 prefusion S trimer. (A) N-terminal segment of S protein. The N-terminus is at residue Gln14 after cleavage of the signal peptide. Cys15 forms a disulfide bond with Cys136. We observed good density for the N-linked glycan at Asn17. (B) A segment immediately downstream of the fusion peptide, while disordered in the stabilized soluble S ectodomain trimer structure, forms a tightly packed structure, designated FPPR for the fusion peptide proximal region, abutting CTD1. The newly identified FPPR structure would clash with CTD1 in the RBD up conformation. Various domains are shown in the color scheme in Fig. 2B. The structure of the soluble S trimer with one RBD in the up conformation (PDB ID: 6vyb) is shown in gray. In the box, a close-up view of the FPPR with adjacent fusion peptide in both surface representation and stick model. (C) The SARS-CoV-2 prefusion S trimer, viewed along the threefold axis, is superposed on the structure of the stabilized soluble S ectodomain trimer in the closed conformation with all three RBDs in the down conformation (PDB ID: 6vxx). While the S2 region is well aligned, there is a significant shift (e.g. ∼10Å between two Ala123 residues) in S1. (D) Impact of the proline mutations introduced at residues 986 and 987 to stabilize the prefusion conformation. K986P mutation removes a salt bridge between Lys986 of one protomer and either Asp426 or Asp427 of another protomer in the trimer interface.

Second, another disulfide containing segment (residues 828-853), immediately downstream of the fusion peptide is also absent from the structures of the soluble ectodomain, but ordered in our new structure (Fig. 3B). We designate it as the fusion-peptide proximal region (FPPR). The FPPR is disordered in both the closed and one RBD-up conformations of the stabilized soluble S trimer. In our new structure of the full-length protein, it packs rather tightly around an internal disulfide bond between Cys840 and Cys851, further reinforced by a salt bridge between Lys835 and Asp848, as well as by an extensive hydrogen bond network. When compared with the RBD-up conformation by superposition of the rest of S2, the FPPR clashes with CTD1, which rotates outwards with RBD in the flipping-up transition. Thus, a structured FPPR, abutting the opposite side of CTD1 from RBD, appears to help clamp down RBD and stabilize the closed conformation of the S trimer. It is not obvious why the FPPR is also not visible in the published, closed S ectodomain structure with all three RBDs in the down conformation^23^. Our structure of the full-length S protein suggests that CTD1 is a structural relay between RBD and FPPR that can sense the displacement on either side. The latter is directly connected to the fusion peptide. Lack of a structured FPPR in the stabilized, soluble S trimer may explain why the RBD-up conformation is readily detected in that preparation. In the 3D classification of our prefusion particles, a large class refined to only a slightly lower resolution (∼3.5Å) with C3 symmetry than did the smaller class that we used for our structure interpretation. That large class did not give any subclasses with RBD flipped up even when reclassified without C3 symmetry (Fig. S5), suggesting that the RBD-up conformation is very rare in our full-length S preparation.

Other major differences between our new structure and the closed conformation of the soluble S trimer stabilized by the proline mutations are large shifts of various sections in S1 when the two structures are aligned by the S2 portion. In Fig. 3C, the three S1 subunits move outwards away from the three-fold axis, up to ∼10Å in peripheral areas (also Fig. S9), suggesting the full length S trimer is more tightly packed among the three protomers than the mutated soluble trimer. When examining the region near the proline mutations between HR1 and CH, we found that the K986P mutation appeared to eliminate a salt bridge between Lys986 in one protomer and either Asp426 or Asp427 in another protomer; thus, the mutation could create a net charge (three for one trimer) inside the trimer interface. This observation may explain why the soluble trimer with the PP mutation has a looser structure than the full-length S with wildtype sequence. Whether this loosening effect by the mutations also led to disordered FPPRs in the closed trimer will require additional experimental evidence. Thus, the proline mutations, originally designed to destabilize the postfusion conformation and strengthen the prefusion structure, may have had some unintended consequences.

### Structure of the postfusion S2 trimer

As expected, 3D reconstruction of the sample from peak 2 yielded a postfusion structure of the S2 trimer, shown in Fig. 4A. The overall architecture of the SARS-CoV-2 S2 in the postfusion conformation is nearly identical to that of the published structure derived from the S2 ectodomain of mouse hepatitis virus (MHV) produced in insect cells (Fig. S1; ref^27^). In the structure, HR1 and CH form an unusually long central three-stranded coiled coil (∼180Å). The connector domain, together with a segment (residues 718-729) in the S1/S2-S2’ fragment, form a three-stranded β sheet, which is invariant between the prefusion and postfusion structures. In the postfusion state, residues 1127-1135 join the connector β sheet to expand it into four strands, while projecting the C-terminal HR2 towards the viral membrane. Another segment (residues 737-769) in the S1/S2-S2’ fragment makes up three helical regions locked by two disulfide bonds that pack against the groove of the CH part of the coiled coil to form a short six helix bundle structure (6HB-1 in Fig. 4B). We do not know whether the S’2 site is cleaved or not in our structure since it is in a disordered region spanning 142 residues (Fig. 4B), as in the MHV S2 structure. Nevertheless, the S1/S2-S2’ fragment is an integral part of the postfusion structure and would not dissociate, regardless of cleavage at the S2’ site. The N-terminal region of HR2 adopts a one-turn helical conformation and also packs against the groove of the HR1 coiled-coil; the C-terminal region of HR2 forms a longer helix that makes up the second six-helix bundle structure with the rest of the HR1 coiled-coil (6HB-2 in Fig. 4B). Thus, the long central coiled-coil is reinforced multiple times along its long axis, making it a very rigid structure, as evident even from 2D class averages of particles in the cryo images (Fig. S7).

**Figure 4.**
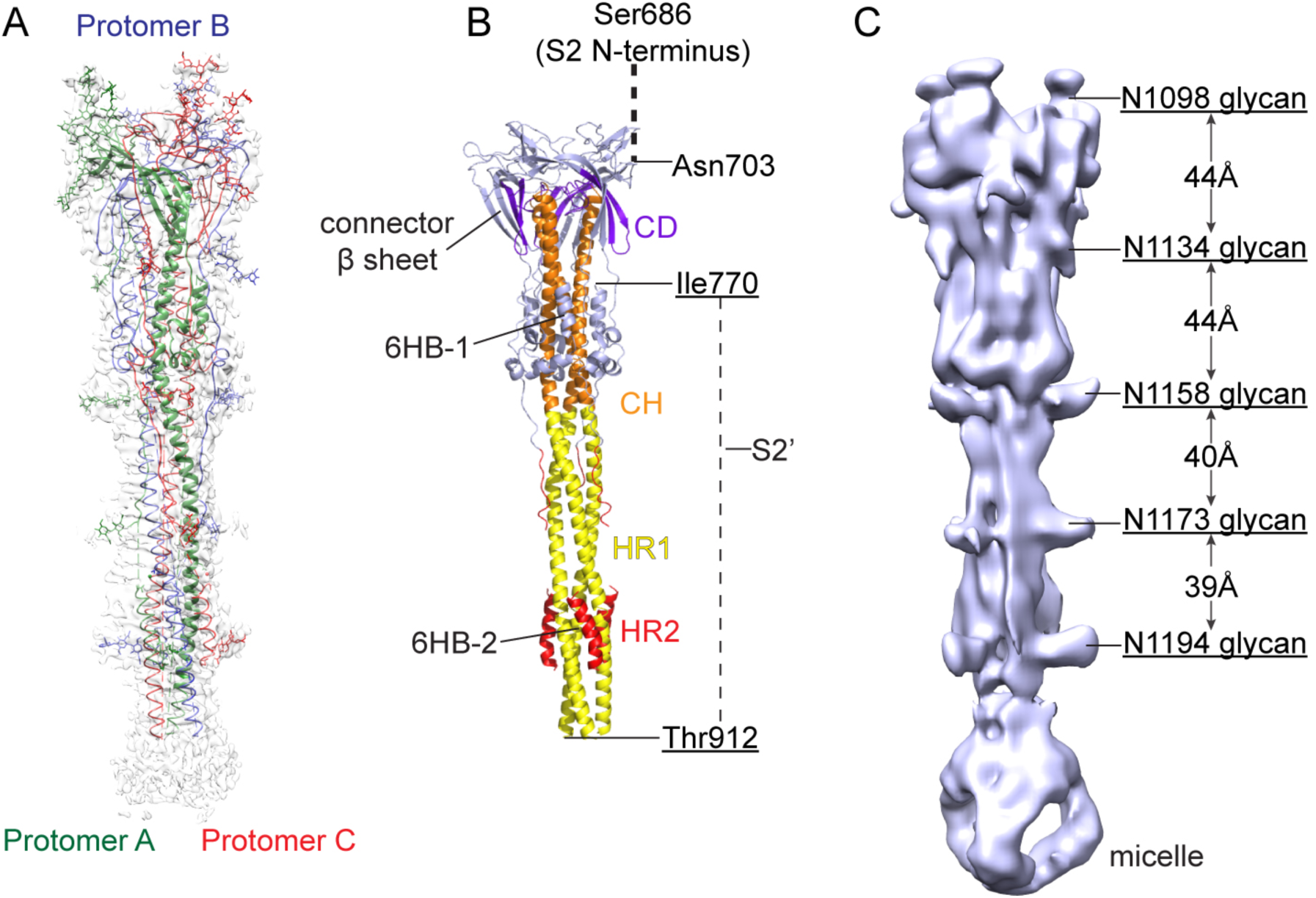
Cryo-EM structure of the SARS-CoV-2 S2 in the postfusion conformation. (A) The structure of the S2 trimer was modeled based on a 3.3Å density map. Three protomers (A, B, C) are colored in green, blue and red, respectively. (B) Overall structure of the S2 trimer in the postfusion conformation shown in ribbon diagram. Various structural components in the color scheme shown in Fig. 1A include HR1, heptad repeat 1; CH, central helix region; CD, connector domain; and HR1, heptad repeat 2. The S2’ cleavage site is in a disordered loop between Ile770 and Thr912. A possible location of the S2 N-terminus (S1/S2 cleavage site) is also indicated. (C) A low resolution map showing the density pattern for 5 N-linked glycans, with almost equal spacing along the long axis.

Another striking feature of the postfusion S2 is its surface decoration by N-linked glycans (Fig. 4C), which are likewise visible from the 2D class averages (Fig. S7). Five glycans at residues Asn1098, Asn1134, Asn1158, Asn1173 and Asn1194 appear to be strategically positioned along the long axis with a regular spacing; four of them are also aligned on the same side of the trimer. If these glycosylation sites are fully occupied by branched sugars, they may shield most surfaces of the postfusion S2 trimer. A similar pattern has been recently described in a paper posted in ChinaXiv (http://www.chinaxiv.org/user/download.htm?id=30394) for a SARS-CoV S2 preparation derived from a soluble S ectodomain construct produced in insect cells and triggered by proteolysis and low pH. Such a decoration seems unrelated to concealment of antigenic surfaces of a functional S trimer, as a postfusion structure has in principle already accomplished its mission.

The fraction from peak 3 contains primarily the dissociated monomeric S1 fragment, which has the smallest size (∼100 kDa) and shows the lowest contrast in cryo grids of the three particle types we describe. We nonetheless extracted ∼40,000 particles and carried out a preliminary 3D reconstruction analysis (Fig. S10), further confirming its identity.

## Discussion

### Architecture of S protein on the surface of SARS-CoV-2 virion

The most unexpected finding from the current study is that the cleaved (S1/S2)_3_ complex dissociates in the absence of ACE2 and that the S2 fragment, along with the S1/S2-S2’ peptide, adopts a postfusion conformation under the mild detergent conditions, suggesting that the kinetic barrier for the conformational transition relevant to viral entry is surprisingly low for this S protein. Whether or not this observation relates directly to efficient membrane fusion or infection, and perhaps effective transmission from human to human leading to the current pandemic, will require further investigation. Nevertheless, it is noteworthy that the postfusion S2 trimer not only has a very stable and rigid structure, but also that it is strategically decorated with N-linked glycans along its long axis with almost even spacing, as if under selective pressure for functions other than as the end-of-mission (postfusion) stage in the membrane fusion process. Although some have suggested that viral fusion proteins may further oligomerize in their postfusion conformation to facilitate fusion pore formation^37^, the protruding surface glycans of the SARS-CoV-2 S2 make this scenario unlikely. A more plausible possibility is a protective role that the S2 postfusion structure could play if it is also present on the surface of an infectious and mature virion. It may induce nonneutralizing antibody responses to evade the host immune system; it may also shield the more vulnerable prefusion S1/S2 trimers under conditions outside the host by decorating the viral surface with interspersed rigid spikes (Fig. 5A). Several recent reports have provided some evidence supporting this possibility. First, EM images of a β-propiolactone inactivated SARS-CoV-2 virus preparation, purified by a potassium tartrate-glycerol density gradient, appeared to have lost all S1 subunits, leaving only the postfusion S2 on the virion surfaces^38^. Likewise, EM images of a β-propiolactone inactivated SARS-CoV-2 virus vaccine candidate (PiCoVacc) also showed needle-like spikes on its surfaces^39^. Second, there are elongated spikes on the surfaces of live viruses in the images released to the news media (http://nmdc.cn/#/nCoV). Third, there are significantly higher antibody binding titers against S2 than those for RBD and S1 in COVID-19 patients^40^, suggesting S2 is more exposed to the host immune system than indicated by the unprotected surfaces on the prefusion structures (ref^22,23^; also Fig. 2). We therefore suggest that postfusion S2 trimers may have a protective function by constituting part of the crown on the surface of mature and infectious SARS-CoV-2 virion (Fig. 5). The postfusion S2 spikes are probably formed after spontaneous dissociation of S1, totally independent of the target cells.

**Figure 5.**
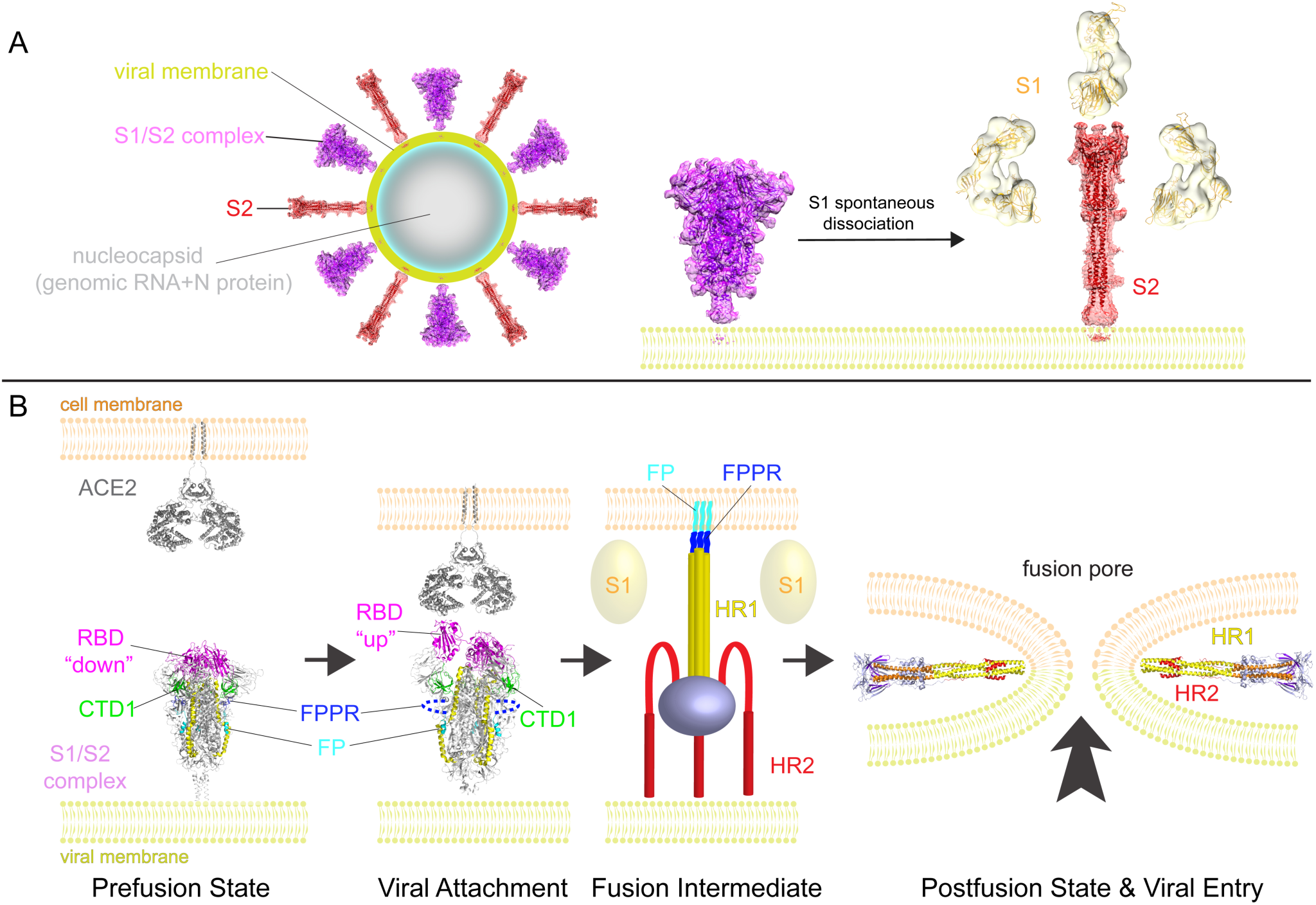
A model for structural rearrangements of SARS-Cov-2 S protein. (A) Structural changes independent of a target cell. We suggest that both the prefusion and postfusion spikes are present on the surface of mature virion and the ratio between them may vary (diagram of virion). The postfusion spikes on the virion are formed by S2 after S1 dissociates in the absence of ACE2. (B) ACE2-dependent structural rearrangements. Structural transition from the prefusion to postfusion conformation inducing membrane fusion likely proceeds stepwise as follows: 1) FPPR clamps down RBD through CTD1 in the prefusion S trimer, but it occasionally flips out of position and allows an RBD to sample the up conformation. 2) RBD binding to ACE2 creates a flexible FPPR that enables exposure of the S2’ cleavage site immediately upstream of the adjacent fusion peptide (FP). Cleavage at the S2’ site, and perhaps also the S1/S2 site, releases the structural constraints on the fusion peptide and initiates a cascade of refolding events in S2, probably accompanied by complete dissociation of S1. 3) Formation of the long central three-stranded coiled-coil and folding back of HR2. 4) Formation of the postfusion structure of S2 that brings the two membranes together, facilitating formation of a fusion pore and viral entry.

### Membrane fusion

Our current study has identified a structure near the fusion peptide – the fusion peptide proximal region (FPPR), that may play a critical role in the fusogenic structural rearrangements of S protein. There appears to be crosstalk between the RBD and the FPPR, mediated by CTD1, as a structured FPPR clamps down RBD while an RBD-up conformation disorders the FPPR by pushing out of its position when the RBD is down. Moreover, the FPPR is also close to the S1/S2 boundary and the S2’ cleavage site, and thus might be the center of activities relevant to conformational changes in S. We do not know the sequence of events, however, nor are we sure about the distinct roles of cleavages at the S1/S2 and S’2 sites. One possibility is that the FPPRs clamp down the three RBDs in the prefusion S trimer, but one occasionally flips out of position due to intrinsic protein dynamics, allowing RBDs to sample the up conformation. A fluctuation of this kind would loosen the entire S trimer, as observed in modified soluble S trimer constructs^22,23^. Once an RBD is fixed in the up position by binding to ACE2 on the surface of a target cell, a flexible FPPR may enable exposure of the S2’ cleavage site immediately upstream of the adjacent fusion peptide. Cleavage at the S2’ site releases the structural constraints on the fusion peptide, which may initiate a cascade of refolding events in S2, including formation of the long central three stranded coiled-coil, folding back of HR2 and ultimately membrane fusion. Cleavage at the S1/S2 site allows complete dissociation of S1, which may also facilitate S2 refolding.

Puzzles regarding membrane fusion remain, as the regions near the viral membrane are still not visible in the reconstructions, even in our structures obtained using a full-length S construct. Yet these regions all play critical structural and functional roles. For example, the conserved hydrophobic region immediately preceding the TM domain, and possibly the TM itself, have been shown to be crucial for S protein trimerization and membrane fusion^29^. The cytoplasmic tail, containing a palmitoylated cysteine-rich region, is believed to be involved in viral assembly and cell-cell fusion^30-33^. The HR2 is unlikely to be disordered on the surface of virion since most spikes appear to stand straight relative to the viral membrane from the EM images of the virus. Whether other viral proteins, such M protein, may help stabilize the spike by interacting with the HR2 remains an interesting question. Thus, we still need a high-resolution structure of an intact S protein in the context of the membrane and other viral components to answer the various open questions.

### Considerations for vaccine development

A safe and effective vaccine is the only medical option to reduce or eliminate the constant threat posed by SARS-CoV-2. The first round of vaccine candidates with various forms of the spike (S) protein of the virus, largely modeled on those designed against SARS-CoV and MERS-CoV^1,16^, are passing rapidly through preclinical studies in animal models and clinical trials in humans. Our study raises several potential concerns about the current vaccine strategies. First, vaccines using the full-length wildtype sequence of S protein may produce the various forms in vivo that we have observed here. The postfusion conformations could expose immunodominant, nonneutralizing epitopes that distract the host immune system, as documented for other viruses, such as HIV-1 and RSV^41,42^. Second, the approach to stabilize the prefusion conformation by introducing proline mutations at residues 986 and 987 may not be optimal, as the K986P mutation may break a salt bridge between protomers that contributes to the trimer stability. The resulting S trimer structure with a relaxed apex may induce antibodies that could not efficiently recognize S trimer spikes on the virus. Third, in light of the possibility that the postfusion S2 is present on infectious virions, vaccines using β-propiolactone inactivated viruses may require additional quality control tests. Although the PiCoVacc appears to provide protection against challenges in nonhuman primates after three immunizations^39^, it is unclear how to minimize the number of the postfusion S2 trimers to avoid batch variations, because another preparation of β-propiolactone inactivated viruses has only the postfusion S2 on the surfaces, which may not elicit sufficient protective antibody responses. Furthermore, sobering lessons from development of SARS-CoV vaccines suggest that some S protein-based immunogens induce harmful immune responses to liver or lung in animal models^43,44^, as well as antibody-dependent enhancement (ADE) of infectivity^45^. It will be critical to define structural determinants that distinguish the ineffective or deleterious responses from the protective responses, to refine next-generation vaccine candidates. Refined immunogens will be particularly critical if SARS-CoV-2 becomes seasonal and returns with antigenic drift, as do influenza viruses^46^.

### Implications for therapeutics

Although virus-encoded enzymes (e.g., RNA-dependent RNA polymerase and proteases) are excellent therapeutic targets, fusion inhibitors that block the conformational changes of S protein may also be promising drug candidates. Fusion inhibitors may even be advantageous because, like antibodies, they do not need to cross cell membrane to reach their target. Moreover, S protein has no obvious cellular homologs and functions by a unique mechanism, and it is thus a more probable target for inhibitors with high specificity but fewer side effects than inhibitors of the viral enzymes. For example, we have recently identified several small-molecule fusion inhibitors, guided by a neutralizing antibody, against HIV-1 envelope spike^47^. These compounds specifically inhibit the envelope-mediated membrane fusion by blocking CD4-induced conformational changes. Because they target a highly conserved site, they also inhibit entry of related viruses, such as HIV-2 and SIV, raising the possibility that similar broad fusion inhibitors can be developed against a diverse set of coronaviruses.

The spread of SARS-CoV-2 has changed our perception of viruses previously thought to be containable. Structural analysis, accelerated by technological advances such as cryo-EM, enable rapid generation of in-depth knowledge of the molecular characteristics of this virus. Our work, which represents one of many complementary studies, may guide our responses to the spread of SARS-CoV-2 in a more rational way than would have been possible even a few months ago.

## Acknowledgments

We thank M. Liao for generous advice, SBGrid team for technical support, and S. Harrison, M. Liao, A. Carfi and D. Barouch for critical reading of the manuscript. Negative stain and cryo-EM data were collected at the Molecular Electron Microscopy Suite and the Harvard Cryo-EM Center for Structural Biology, respectively, at Harvard Medical School. This work was supported by NIH grants AI147884 (to B.C.), 3R01AI147884 (to B.C), AI141002 (to B.C.), AI127193 (to B.C. and James Chou), as well as a COVID19 Award by Massachusetts Consortium on Pathogen Readiness (MassCPR; to B.C.).

## Author Contribution

B.C. and Y.C. conceived the project. Y.C. and H.P. expressed and purified the full-length S protein. T.X. expressed and purified soluble ACE2, performed SPR and cell-cell fusion experiments. Y.C. and J.Z. performed negative stain EM analysis. J.Z. prepared cryo grids and performed EM data collection with contributions from S.S. and R.M.W.. J.Z. processed the cryo-EM data, built and refined the atomic models for the prefusion S trimer and the postfusion S2 trimer. Y.C. processed the S1 data. S.R. contributed to data processing for S1 and provided computational support. S.R.V. contributed to cell culture and protein production. All authors analyzed the data. B.C., Y.C., J.Z. and T.X. wrote the manuscript with input from all other authors.

## Methods

### Expression constructs

Genes of a full-length spike (S) protein (residue 1-1273) of SARS-CoV-2 and the ectodomain of human angiotensin converting enzyme 2 (ACE2; residue 1-615) were synthesized by GenScript (Piscataway, NJ). The S gene was fused with a C-terminal twin Strep tag [(GGGGS)_2_WSHPQFEK(GGGGS)_2_WSHPQFEK)] and cloned into a mammalian cell expression vector pCMV-IRES-puro (Codex BioSolutions, Inc, Gaithersburg, MD). The ACE2 gene added a C-terminal 6xhistidine tag and cloned into the same expression vector.

### Expression and purification of recombinant proteins

Expi293F cells (Thermo Fisher Scientific) were transiently transfected with the S protein expression construct using polyethylenimine (PEI; Sigma-Aldrich) in FreeStyleMedia (Gibco) containing 1% Pen Strep (Gibco). Protein expression was confirmed by western blot (see below). To purify the full-length S protein, the transfected cells were harvested at a density of ∼4-5×10^6^/ml by centrifugation (1,000 xg at 4°C for 30 min). Cell pellet was washed with PBS, resuspended in a lysis buffer containing Buffer A (100 mM Tris-HCl, pH 8.0, 150 mM NaCl, 1 mM EDTA) and 1% NP-40 (w/v; AmericanBio, Natick, MA), EDTA-free complete protease inhibitor cocktail (Roche, Basel, Switzerland), and incubated at 4°C for one hour. After clarifying spin (30,000 xg at 4°C for 60 min), the supernatant was loaded on a strep-tactin (IBA Lifesciences, Germany) column equilibrated with the lysis buffer. The column was then washed with 50 column volumes of Buffer A and 0.3% NP-40, followed by additional washes with 50 column volumes of Buffer A and 0.1% NP-40, and with 50 column volumes of Buffer A and 0.02% NP-40. The S protein was eluted by Buffer A containing 0.02% NP-40 and 5 mM desthiobiotin. Elution fractions were analyzed by SDS-PAGE (Bio-Rad), and those containing S protein were pooled and concentrated. The protein was further purified by gel filtration chromatography on a Superose 6 10/300 column (GE Healthcare) in a buffer containing 25 mM Tris-HCl, pH 7.5, 150 mM NaCl, 0.02% NP-40.

HEK293T cells transfected with the his-tagged ACE2 expression construct were grown in 250 ml roller bottles with FreeStyleMedia containing 1% Pen Strep. The protein was purified by affinity chromatography using Ni-NTA agarose (Qiagen, Hilden, Germany), followed by gel filtration chromatography, as described previously^48,49^. The peak fractions were pooled and concentrated to 10 mg/ml using a 10 kDa MWCO Millipore filter (MilliporeSigma, Burlington, MA).

### Western blot

Western blot was performed either using an anti-strep tag antibody or anti-SARA-COV-2 S antibody following a protocol described previously^50^. Briefly, full-length S protein samples were resolved in 4-15% Mini-Protean TGX gel (Bio-Rad) and transferred onto PVDF membranes (Millipore, Billerica, MA) by an Iblot2 (Life Technologies). Membranes were blocked with 5% skimmed milk in PBS for 1 hour and incubated either with anti-strep tag antibody (IBA Lifesciences) or anti-SARS-COV-2 polyclone antibody (Sino Biological Inc.) for another hour at room temperature. Alkaline phosphatase conjugated anti-Rabbit IgG (1:5000) (Sigma) was used as a secondary antibody. Proteins were visualized using one-step NBT/BCIP substrates (Promega).

### Negative stain EM

To prepare grids, 3 µl of freshly purified full-length S protein was adsorbed to a glow-discharged carbon-coated copper grid (Electron Microscopy Sciences), washed with deionized water, and stained with freshly prepared 1.5% uranyl formate. Images were recorded at room temperature at a magnification of 67,000x and a defocus value of 2.5 µm following low-dose procedures, using a Tecnai T12 electron microscope (Thermo Fisher Scientific) equipped with a Gatan UltraScan 895 4k CCD camera and operated at a voltage of 120 keV. Particles were auto-picked, and 2D class averages generated using RELION software 3.0.8.

### Cell-cell fusion assay

The cell-cell fusion assay, based on the α-complementation of E. coli β-galactosidase, was conducted to quantify the fusion activity mediated by SARS-CoV2 S protein. Varying amount of the full-length SARS-CoV2 S protein expression construct (0.1-10 µg) and the α fragment of β-galactosidase construct (10 µg), or the full-length ACE2 expression construct (10 µg) together with the ω fragment of β-galactosidase construct (10 µg), were mixed with DMEM medium (Gibco) containing PEI (80 µg) and added to HEK293T cells. Following a 5-hr incubation at 37°C, the medium was aspirated and replaced with complete DMEM (1% Pen Strep, 1% GlutaMax and 10% FBS). After an additional 19-hour incubation at 37°C, the cells were detached using PBS with 5 mM EDTA and resuspended in complete DMEM. 50 µl S-expressing cells (1.0×10^6^ cells/ml) were mixed with 50 µl ACE2-expressing cells (1.0×10^6^ cells/ml) to allow cell-cell fusion proceed at 37°C for 2 hours. Cell-cell fusion activity was quantified using a chemiluminescent assay system, Gal-Screen (Applied Biosystems, Foster City, CA), following the standard protocol recommended by the manufacturer. The substrate was added to the mixture of the cells and allowed to react for 90 minutes in dark at room temperature. The luminescence signal was recorded with a Synergy Neo plate reader (Biotek).

### Surface Plasmon Resonance (SPR) analysis

All experiments were performed with a Biacore 3000 system (GE Healthcare) at 25°C in HBS buffer (10 mM HEPES, pH 7.0, 150 mM NaCl, 3 mM EDTA, 0.005% P20 surfactant). Protein immobilization to CM5 chips was performed following the standard amine coupling procedure as recommended by the manufacturer. Various forms of SARS-CoV2 S protein were immobilized at a level of ∼3000 RU. Sensorgrams were recorded by passing various concentrations (15.6-250 nM) of soluble ACE2 over the S-immobilized surface at a flow rate of 40 µl/min with a 4-min association phase followed by a 10-min dissociation phase. The surface was regenerated by dissociation in the running buffer for another 2 minutes. Identical injections over blank surfaces were subtracted from the data for kinetic analysis. Binding kinetics was analyzed by BIAevaluation software using a 1:1 Langmuir binding model. All injections were carried out in duplicate and gave essentially identical results.

### Cryo-EM sample preparation and data collection

To prepare cryo grids, 3.5 µl of each freshly purified fractions from peak 1-3 (see Fig. 1B) at ∼0.3 mg/ml was applied to a 1.2/1.3 Quantifoil grid with continuous carbon support (Quantifoil Micro Tools GmbH), which had been glow discharged for 60 s at 15 mA. Grids were immediately plunge-frozen in liquid ethane using a Vitrobot Mark IV (Thermo Fisher Scientific) with a blotting time of 4 s. The grids were first screened for ice thickness and particle distribution using a Talos Arctica transmission electron microscope (Thermo Fisher Scientific), operated at 200 keV and equipped with a K3 direct electron detector (Gatan), at the Harvard cryo-EM center for Structural Biology. For data collection, images were acquired with selected grids using a Titan Krios transmission electron microscope (Thermo Fisher Scientific) operated at 300 keV with a BioQuantum GIF/K3 direct electron detector. Automated data collection was carried out using SerialEM version65 at a magnification of 105,000× and the K3 detector in counting mode (pixel size, 0.825 Å) at a dose rate of ∼1.1 electrons per physical pixels per second. Each movie had a total accumulated exposure of ∼50 e/Å2 fractionated in 50 frames of 200 ms. A dataset for each sample was acquired using a defocus range of 1.5-2.6 µm.

### Image processing and 3D reconstructions

Drift correction for cryo-EM images was performed using MotionCor2^51^, and contrast transfer function (CTF) was estimated by CTFFIND4^52^ using motion-corrected sums without dose-weighting. Motion corrected sums with dose-weighting were used for all other image processing. RELION3.0.8 was used for particle picking, 2D classification, 3D classification and refinement procedure. Approximately 3,000 particles were manually picked for each protein sample and subjected to 2D classification to generate the templates for automatic particle picking. For the peak 1 sample, after manual inspection of auto-picked particles, a total of 1,069,976 particles were extracted from 19,013 images. The selected particles were subjected to 2D classification, giving a total of 338,930 good particles. The low-resolution negative-stain reconstruction of the sample was low-pass-filtered to 40Å as an initial model for 3D classification with C3 symmetry. Two major classes with 42,638 and 288,090 particles, respectively, showed clear structural features were subjected to 3D refinement with C3 symmetry using an overall mask. The smaller class gave a reconstruction at 3.4Å resolution while the large class led to a map at 3.5 Å. The particles from the smaller class were subjected to CTF refinement and Bayesian polishing, followed by another round of 3D refinement, resulting in a reconstruction at 3.1Å resolution, which was used for structural interpretation. The particles from the larger class were reclassified without C3 symmetry, giving only one major class with no significant differences in the map as compared to the previous round. One possibility that the larger class did not refine better than the smaller class might be due to slight distortion of the particles when landed on the carbon surface.

For images of the sample from peak 2, after manual inspection of auto-picked particles, a total of 2,138,443 particles were extracted from 17,909 images. The selected particles were subjected to 2D classification, leading to a total of 1,007,156 good particles. 2D class averages of these particles were then used to create a 3D initial model in RELION. After two rounds of 3D classification, a total of 443,148 particles were subjected to 3D refinement, yielding a reconstruction at 4Å resolution. Two additional rounds of 3D classification without alignment using an overall mask gave a class with a total of 196,506 particles showing clear structural features. These particles were subjected to 3D refinement with overall mask, yielding a reconstruction at 3.3Å resolution. Local refinement with top and bottom region masks improved the local resolution to 3.2Å for the top and 3.7Å resolution for the bottom, respectively.

Reported resolutions are based on the gold-standard Fourier shell correlation (FSC) using the 0.143 criterion. All density maps were corrected from the modulation transfer function of the K3 detector and then sharpened by applying a temperature factor that was estimated using post-processing in RELION. Local resolution was determined using RELION with half-reconstructions as input maps.

For the S1 fragment data set, CrYOLO^53^ was used for particle picking, RELION 3.0.8 was used for 2D classification, 3D classification and refinement. For a preliminary analysis, a total of 40,000 particles were extracted from 2,264 selected images. They were subjected to 2D classification, giving a total of 26,533 particles, which were first used to generate an initial 3D model in RELION. After 3D classification, a class of 6,975 particles showing a V-shaped density was refined with an overall mask, giving a reconstruction at 12.5Å resolution, as reported by RELION. The resolution may not be accurate because the small number of particles used. The S1 model derived from our prefusion structure was fit into the map by Chimera^54^ and Coot^55^. Structural biology applications used in this project were compiled and configured by SBGrid^56^

### Model building

The initial templates for model building used the stabilized SARS-CoV-2 S ectodomain trimer structure (PDB ID 6vxx) for the prefusion conformation, and a homology model, calculated by I-TASSER, of S2 postfusion trimer structure from mouse hepatitis virus (MHV) (PDB ID: 6b3o) for the postfusion confirmation. Several rounds of manual building were performed in Coot. For the prefusion structure, the model was refined in Phenix^57^ against the 3.1Å cryo-EM map. For the postfusion structure, the model was first refined in Phenix against two locally refined 3.2Å and 3.7Å maps, and then refined against the overall 3.3Å map. The refinement statistics are summarized in Extended Data Table1.

